# Co-occurrence networks from wetland bird surveys predict H5N1 genetic similarity between wild birds with strong effects of time, space and viral clustering

**DOI:** 10.1101/2025.06.17.659947

**Authors:** Jamie Dunning, Anna Gamża, Josh A. Firth, Adham Ashton-Butt, William T. Harvey, Samantha J. Lycett, Rowland R. Kao, Ian H. Brown, Alastair I. Ward

## Abstract

The emergence of zoonotic and epizootic diseases has had devastating consequences for human and animal health, including wildlife conservation. Yet, surveillance of multi-host disease systems is particularly challenging due to complex transmission pathways across many species. Social network analysis has been applied to simple transmission systems, but empirical applications to wild, multi-species systems are scarce. Here, we combined high pathogenicity avian influenza (HPAI) viral genomes, a zoonotic virus of pandemic potential, with a large citizen-science database of wild bird co-occurrence to test how multi-species social network structure predicts transmission dynamics. We linked viral genetic distance, 20,103 pairwise comparisons between 214 unique genomes from 172 dyads of 20 host species, to co-occurrence network metrics for those species. Both relative species association and raw co-occurrence frequency predicted lower maximum viral genetic divergence, more similar viruses between more associated species, beyond what would be expected through random mixing and independently of sequencing effort. Time and space between samples were also strong predictors of genetic similarity. Our results suggest that network models can be used to detect pathogen transmission through communities of wild birds, offering real prospects for wildlife disease surveillance and prediction.

## Introduction

Emerging zoonotic and epizootic pathogens that spill over from wildlife are a persistent, and often intractable, threat to human and animal health (Kruse 2004; Jones et al. 2008) with far-reaching consequences for global economic and food security (Bernstein et al. 2022). Many such diseases, including Ebola and SARS-CoV-2, have originated from multi-species groups of wild animals, often without detection in those populations prior to disease emergence in humans or livestock (Marí et al. 2015; Holmes 2022; Xie et al. 2023). This is particularly true in multi-host-pathogen systems like high pathogenicity avian influenza viruses (HPAI) in wild birds, where ecological and behavioural diversity across many host species gives rise to complex transmission pathways (Bansal et al. 2007; Buhnerkempe et al. 2015; Dunning et al. 2025), presenting a challenge for proactive surveillance. Until recently, network models of disease systems have adopted single-species, trait- or behaviour-based approaches (Vickers et al. 2024; Gamża et al. 2024; Yin et al. 2023; Luis et al. 2015; Olival et al. 2017; McDuie et al. 2024). Understanding how multi-species interactions in the wild shape disease spread therefore represents an important knowledge gap for informing zoonotic and epizootic disease surveillance (Buhnerkempe et al. 2015; Sah et al. 2018; Albery et al. 2021; Dunning et al. 2025).

In communities of non-human animals, patterns of disease transmission are highly variable and cannot be solely explained by pathogen virulence. Instead, just as human mobility and co-occurrence in space, influences the trajectory of an epidemic (Firth et al. 2020; Kostandova et al. 2024), animal social network structure, patterns of association between individuals and species shaped by the social (Bansal et al. 2007; Sah et al. 2017; Albery et al. 2021) and physical environment (Carlson et al. 2022; King et al. 2023), influence epizootic disease propagation. The mechanism by which these host traits, or environmental components influence disease transmission is often by shaping social network structures - forcing potential hosts together or driving them apart. This spatial and temporal clustering either facilitates or constrains non-vector borne pathogen transmission through social links (Hamede et al. 2009). For example, during the HPAI outbreak of 2021/22, H5N1 clade 2.3.4.4b began infecting European seabirds. The subsequent redrawing of host transmission networks through newly accessible flyways, had devastating consequences for the conservation of wild birds (Falchieri et al. 2022; Tremlett et al. 2025) driven, in part, through co-occurrence inside colonial breeding colonies (Matthiopoulos et al. 2026). These interactions also likely drove incursion of HPAI to new continents (Kandeil et al. 2023) and novel mammalian hosts (Sidik 2023; Plaza et al. 2024; Sah et al. 2024). Recent work has underlined the role of specific host traits on social network structure through this outbreak (Wang et al. 2026), yet, the ways in which species co-occurrence drives infection at the landscape scale remains poorly understood. Repeated incursions of different viral genotypes may also structure variation outside of transmission in the landscape (Xie et al. 2023; Harvey et al. 2025)-that is, two species may carry highly divergent viruses because they were infected through different incursion events.

Past evaluations of complex, multi-host network disease dynamics have been largely reliant on theoretical models (Albery et al. 2020; McDuie et al. 2024), while surveillance efforts are limited by reactive sample collection, taxonomic sampling biases, and processing constraints (Wade et al. 2022; Atkinson et al. 2025). Complex network approaches offer promise (Dunning et al. 2025) but have so far only been applied to simpler transmission systems in relatively few species (Hamede et al. 2009; Gómez et al. 2013; Wilson-Aggarwal et al. 2019; Proboste et al. 2024). While Dunning et al. (2025) suggested a theoretical framework for HPAI surveillance, empirical evidence from a wild, multi-species system is lacking. Here, we used data from HPAI outbreaks in Great Britain (GB), a serious public health threat with trans-boundary transmission between hundreds of interacting hosts and multiple transmission pathways (Xie et al. 2023; Reid et al. 2024) to test the relationship between multi-species social links and disease transmission. We defined social links broadly, using co-occurrence to encompass shared space and time, and the potential for interaction, the gambit-of-the-group (Franks et al. 2010). We combined social network analysis with viral phylogenetics and Bayesian multilevel modelling to test our prediction that social links predict shorter distance between viral genomes (as a proxy for transmission probability) in multi-species networks of wild birds. To capture potential transmission pathways, we used fine-scale spatiotemporal co-occurrence between commonly occurring waterbird species, derived from national monitoring data. Co-occurrence networks derived from these data likely operate at coarser spatial and temporal resolution than rare transmission events, and their ability to detect real-world signals is an open question. Previous phylodynamic studies have matched H5N1 infection in poultry with identical genotypes in nearby wetland sediments (Giacinti et al. 2024), implying landscape transmission without identified pathways. Our results provide an empirical basis for using co-occurrence networks for wildlife disease surveillance, when applied to multi-species systems at the landscape scale.

## Materials and Methods

To test the hypothesis that host species’ social links predict viral genetic distance across multi-species networks of wild birds, we combined social network analysis with phylodynamic and Bayesian mixed modelling approaches. We also tested if our results differed from those derived from an expected network, without social structure, by running two distinct permutation tests that mirrored our observed data models.

### Survey effort and Networks

To build species co-occurrence networks, we accessed six years of data from the British Trust for Ornithology Wetland Bird Survey (WeBS; Woodward et al. 2024), 2018 - 2023 for two overlapping regions of East Yorkshire, Great Britain, each defined by a 50km circular buffer around two recurrent HPAI outbreak sites. In total 3,722 individual surveys, including 159 species. WeBS follows a standardized survey methodology and aligns surveys with high tides across the year to maximise detection probability of priority wetland birds (Woodward et al. 2024). We treated each survey, a single site for each month of a single WeBS year, as an event and built a single network for each year, where the nodes represented species recorded within an event and their interconnecting edges represented a shared event between a species pair. Combined across years, the global network comprised 8,162 edges between 159 species (for a summary see Figure 1; Table 1a–c).

**Table 1a-c.** Summary statistics for social networks within (a) and between (b) years, and node (viral host species) attributions (c). 1a shows the number of species involved in each network, edges between species pairs representing co-occurrence within a recording event, and the number of recording events. A recording event represents a single survey visit in time and space within the study area. 1b shows between-year network similarity, where the upper-right matrix shows Jaccard similarity (J), the proportion of shared dyads between years irrespective of edge weight; and the lower-left matrix shows Spearman rank correlation coefficients (ρ) between dyadic edge weights for dyads present in both years. 1c shows the 20 host species included in both the viral phylogeny and co-occurrence network that make up those used in our models. Here, host is the species from which virus was sequenced, n events shows the number of events that host was recorded in, degree is the number of distinct connections (overlaps) in the social network, and strength is the sum of edge weights across those connections. Dyadic weight and dyadic SRI give the mean and sd of those association metrics across each species’ dyads in the final model dataset, and n Seq denotes the number of sequences accessed for each host species.

| a. Network structure |  |  |  | b. Network similarity between years |  |  |  |  |  |  |
| --- | --- | --- | --- | --- | --- | --- | --- | --- | --- | --- |
| Year | Species | Edges | Events | J →<br>ρ ↓ | 2018 | 2019 | 2020 | 2021 | 2022 | 2023 |
| 2018 | 115 | 4132 | 354 | 2018 |  | 0.67784878 | 0.65510949 | 0.56959379 | 0.5687052 | 0.57280494 |
| 2019 | 131 | 5321 | 729 | 2019 | 0.86258974 | - | 0.71297379 | 0.64242594 | 0.67200933 | 0.61539724 |
| 2020 | 127 | 4938 | 717 | 2020 | 0.81935908 | 0.89806925 | - | 0.66118614 | 0.63261702 | 0.65320748 |
| 2021 | 134 | 6378 | 782 | 2021 | 0.76063755 | 0.84100878 | 0.83690949 | - | 0.73934768 | 0.62826834 |
| 2022 | 137 | 6154 | 748 | 2022 | 0.75728319 | 0.84711208 | 0.83417479 | 0.93653696 | - | 0.63485986 |
| 2023 | 118 | 4520 | 392 | 2023 | 0.67831702 | 0.76854005 | 0.78439467 | 0.89199321 | 0.90917122 | - |

| c. Node attributes |  |  |  |  |  |  |  |  |
| --- | --- | --- | --- | --- | --- | --- | --- | --- |
| Host | n Events | Degree | Strength | Mean Weight | SD Weight | Mean SRI | SD SRI | n Seq |
| *Anas platyrhynchos* | 3442 | 154 | 45728 | 132.17 | 145.46 | 0.23 | 0.23 | 3 |
| *Cygnus olor* | 2448 | 150 | 40055 | 115.62 | 130.85 | 0.24 | 0.23 | 47 |
| *Fulica atra* | 2386 | 150 | 39239 | 112.93 | 128.37 | 0.24 | 0.23 | 1 |
| *Anser anser* | 2079 | 153 | 35327 | 106.6 | 115.45 | 0.24 | 0.21 | 29 |
| *Branta canadensis* | 2001 | 149 | 33712 | 103.05 | 112.81 | 0.24 | 0.21 | 18 |
| *Vanellus vanellus* | 1043 | 146 | 23878 | 65.83 | 61.28 | 0.21 | 0.11 | 1 |
| *Larus argentatus* | 915 | 152 | 18654 | 55.39 | 50.83 | 0.19 | 0.1 | 12 |
| *Larus canus* | 780 | 148 | 17206 | 49.83 | 43.84 | 0.18 | 0.1 | 3 |
| *Buteo buteo* | 503 | 141 | 13214 | 38.48 | 27.18 | 0.19 | 0.11 | 20 |
| *Falco tinnunculus* | 414 | 143 | 11999 | 33.65 | 22.68 | 0.19 | 0.1 | 2 |
| *Accipiter nisus* | 279 | 136 | 8447 | 24.36 | 17.28 | 0.15 | 0.08 | 4 |
| *Corvus corone* | 316 | 132 | 7463 | 22.96 | 23.56 | 0.12 | 0.07 | 1 |
| *Anser brachyrhynchus* | 246 | 136 | 7054 | 19.45 | 13.96 | 0.12 | 0.06 | 14 |
| *Sterna hirundo* | 255 | 122 | 6093 | 17.99 | 16.03 | 0.09 | 0.04 | 12 |
| *Branta leucopsis* | 188 | 134 | 5831 | 16.02 | 12.16 | 0.1 | 0.05 | 26 |
| *Cygnus cygnus* | 154 | 119 | 4294 | 12.6 | 8.84 | 0.08 | 0.04 | 7 |
| *Falco peregrinus* | 106 | 140 | 3949 | 10.54 | 8.03 | 0.09 | 0.07 | 8 |
| *Hydrocoloeus minutus* | 39 | 111 | 1849 | 5.3 | 3.09 | 0.05 | 0.04 | 1 |
| *Morus bassanus* | 2 | 76 | 92 | 1 | 0 | 0.02 | 0.01 | 4 |
| *Fulmarus glacialis* | 1 | 24 | 24 | 1 | 0 | 0.01 | 0.02 | 1 |

**Figure 1.**
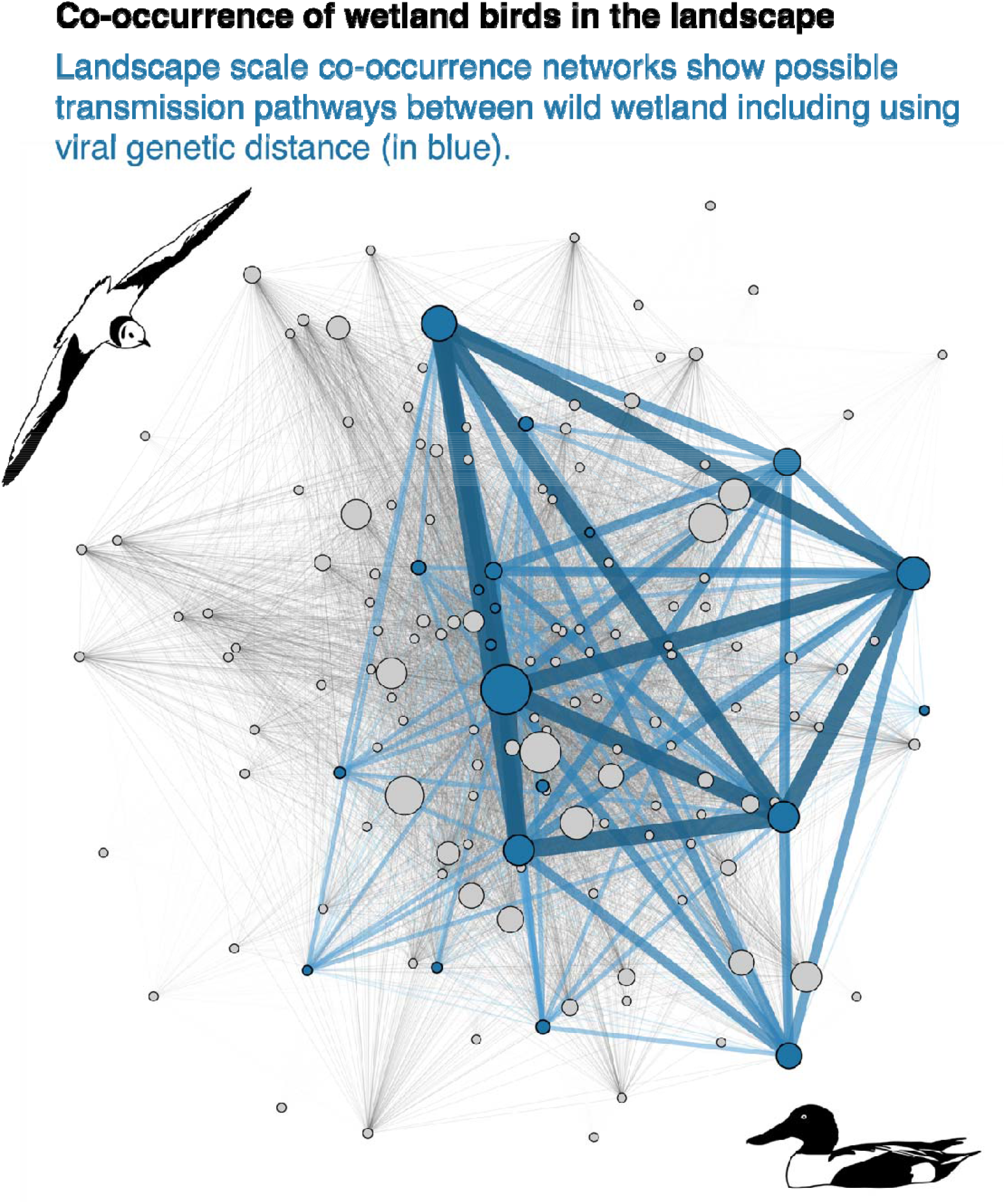
Co-occurrence network of wetland birds and viral sampling coverage. Of the 159 species in the co-occurrence network, 20 were also associated with a genome for high pathogenicity avian influenza clade 2.3.4.4b, of which we had access to 214 unique sequences. The colour of the nodes and edges denote that a species pair had a viral genome available (blue), or not, (grey). The node size denotes normalised betweenness centrality, where the largest nodes are those most central to network structure, in this case Mute Swan *Cygnus olor*, and 0, on none of the shortest paths (here including Northern Lapwing *Vanellus vanellus*, Little egret *Egretta garzetta*, and Northern pintail *Anas acuta*). Edge width denotes extent of co-occurrence, where thicker edges link species pairs more often recorded together.

From these networks we extracted two established dyadic (pairwise - between two species) measures of association. To measure the extent to which two species co-occur in the landscape, and therefore, may be expected to share viruses more easily, we used pairwise edge weight, or simple connection strength, which represented the absolute frequency of co-occurrence in the network. Then, to capture the extent to which a species pair is associated we also used the Simple Ratio Index (SRI; Farine & Whitehead 2015). SRI represents the connectedness of a species pair relative to the opportunity for association, over-and-above the effect of abundance. A greater SRI is therefore a measure of species clustering probability in the social network. For example, in the WeBS data, the highest SRI values were between relatively scarce species whose habitats overlap, for example, Dipper *Cinclus cinclus* and Grey Wagtail *Motacilla cinerea*, which both use fast flowing upland streams. Although they were both scarce in the landscape, where they were recorded they were almost always recorded together. We calculated each measure within each survey year, retaining one value per species pair per year, before filtering and scaling against our viral data.

### Network similarity

We tested the assumption that our networks were comparable across years - that the dyads that make up the general network structure were similar across our six years of data collection. We did this using two measures designed to compare non-uniform networks, in this case, with variation in the species recorded (the network actors, nodes) between years: 1) Jaccard distance describes the proportion of shared dyads between two networks, regardless of weight; and, 2) a Spearman rank correlation test to measure distance of dyadic edge weights between species pairs present in both networks. In both cases 1 denotes identical network structure, and 0 denotes that a pair of networks are completely different (Table 1b).

### Viral genetic distance

We accessed HPAI viral genomes collected throughout Great Britain from the Global Initiative on Sharing All Influenza Data (GISAID; Shu & McCauley 2017). We associated each sequence with host information and filtered out mammals, domestic birds and taxonomically vague entries (none of which are recorded as part of WeBS surveys), as well as non-H5N1 strains (including, H5N5 and H5N8). We aligned each sequence per segment using MAFFT v7.51149, trimmed them to coding regions, removed insertions present in fewer than 10% of sequences and rejected sequences with nucleotide coverage lower than 80% for all segments. We concatenated segment sequences and compared sequences with all others in the database to quantify a genetic distance between pairwise viral genomes using the Tamura and Nei 1993 model (Tamura & Nei 1993) in the ape package for R (Paradis et al. 2019; R Core Team 2025), which calculates genetic distance as a proportion that differ between each pair of sequence, accounting for difference in rates of transition and transversion mutations. While we limited our viral phylogenetic data to the H5N1 subtype, we identified genomes that had acquired genomic segments from influenza viruses of other low pathogenicity sub-types. Thus, there were two mechanisms of viral evolution influencing genetic distances among these H5N1 viruses: small, incremental changes driven by gradual accumulation of mutations during host-to-host transmission, and greater genetic shifts where reassortment occurs between co-infecting viruses. To test if social network centrality predicted transmission probability, we limited our study to these smaller incremental changes by using a measure of partial genetic distance (hereafter genetic distance), using only non-reassorting haemagglutinin, neuraminidase and matrix protein genome fragments (see Harvey et al. 2025). We recorded the date associated with each sampling event and a location, provided by the Animal and Plant Health Agency, that is, measures of geometric (space between sampling events) and temporal (time between sampling events) distance (Gamża et al. 2024; Harvey et al. 2025). After restricting our data to species present in both the WeBS networks and viral datasets, we used 20,103 pairwise comparisons between 214 unique genomes from 172 dyads of 20 host species– that is, genetic distance, aggregated across the full study period, per species pair, and aligned with network measures computed within years; 913 dyad-years.

### Model fit

To test the relationship between wild bird social network structure and genetic distance between HPAI viruses, and to accommodate the multi-membership sampling within our species dyads, we ran a series of Generalized Linear Mixed Models (GLMMs) using Bayesian Markov Chain Monte Carlo (MCMC) simulations, with the Hamiltonian Monte Carlo (HMC) and No-U-Turn Sampler (NUTS) algorithms, in the brms package in R (R Core Team 2025; Bürkner 2017). To capture rare transmission events between wild birds we used the maximum genetic distance between pairs of sequences (median = 0.015, range = 0.002 – 0.028) as our response variable. We chose maximum as the most divergent, most precautionary, measure of viral genetic distance. Maximum genetic distance correlates strongly with sequencing effort (ρ = 0.77, compared to 0.43 for the mean and −0.45 for the minimum). To account for this confound we refitted both models with log sequencing effort as an additional fixed effect, repeating the rewiring permutation on each (see Results). Sequencing effort was only weakly correlated with network metrics (SRI ρ = 0.057; pairwise weight ρ = 0.105). We repeated all models using mean and minimum genetic distance for comparison, and to assess the contribution of species identity, and we also fitted each model without the multi-membership term.

Otherwise, our models only differed in the first predictor (fixed effect) variable, which was either pairwise weight or SRI. For all models, we used weakly informative priors, and ran each over 4 chains of 4,000 iterations, with a warm-up of 1,000 iterations and a thinning interval of 1. We used Effective Sample Sizes of >1,000 in the posterior and tail distributions to maintain sampling efficiency and a scale reduction factor (Ř) value of 1 to ensure convergence. We also manually checked posterior trace plots to ensure model mixing. Because our study focused on transmission pathways, rather than host susceptibility to infection, we did not constrain the phylogenetic relationship between hosts in our models.

### Weight model

We modelled the pairwise weight of birds present in both the WeBS and the viral phylogenetic data (median = 23, range = 1 – 511), using network measures calculated within survey years. We included the mean of geometric distance (median = 251.9 km, range = 88.6 – 682.9 km) between viral pairwise sample locations and the mean of pairwise temporal distance (time elapsed between sampling events; median = 281 days, range = 8 – 538 days) as fixed effects. We used the mean (rather than the maximum) of geometric- and temporal distance to account for wide variation in sampling and to reduce marginal collinearity. We z-transformed all fixed effects to better align scales between other variables in the model. Because most species appeared across many dyads, we accounted for species specific effects on genetic distance independently of their dyad (i.e. species A + species B, rather than species AB) by including species A and species B as a linked ‘multi-membership’ variable inside our random effects. This approach also avoided over-dispersion.

### SRI model

We repeated the first model, using the same model structure but with SRI (median = 0.116, range = 0.0015 – 0.773) instead of pairwise weight, as a fixed effect. We used the log-odds (using a logit link function) of SRI to better accommodate a linear relationship between SRI and other predictors in the model.

### Permutations

To test if our results differed from random social structure, we ran two permutation tests over 1,000 cycles. These were: 1) Network rewiring, shuffling edges between species pairs while maintaining the degree distribution to control for frequency in the network (Farine 2017); and 2) Node-label randomisation, maintaining network structure while permuting species identity within the viral data, testing if specific dyads carry information beyond species identity alone, for example highly divergent viral lineages. We inferred an effect where the observed estimate fell outside the 95% interval of the permuted distribution.

## Results

We derived a global social network from 3,722 individual wetland bird surveys over six years (for a summary see Table 1a-c), comprising 8,162 dyads between 159 wetland bird species. Of these, 20 species were represented in both the social network and the viral genomic dataset (See Figure1; supplementary Table 1 for GISAID accession numbers and associated metadata). Network structure was consistent across our survey years (Table 1b).

Both relative species association and raw co-occurrence frequency predicted lower maximum viral genetic distance (SRI: estimate = −0.032, 95% CI = −0.054 to −0.010; Weight: estimate = −0.041, 95% CI = −0.065 to −0.017), and more than expected under a network rewiring model (Figure 2; SRI: permuted 95% CI = −0.023 to 0.013, p = 0.005; Weight: permuted 95% CI = −0.016 to 0.017, p = 0.001); but neither did under a node-label randomisation (Figure 2; SRI: p = 0.221; Weight: p = 0.153). The permuted distribution of the node-label randomisation had a standard deviation of 0.031–0.034, exceeding estimated effect sizes.

**Figure 2.**
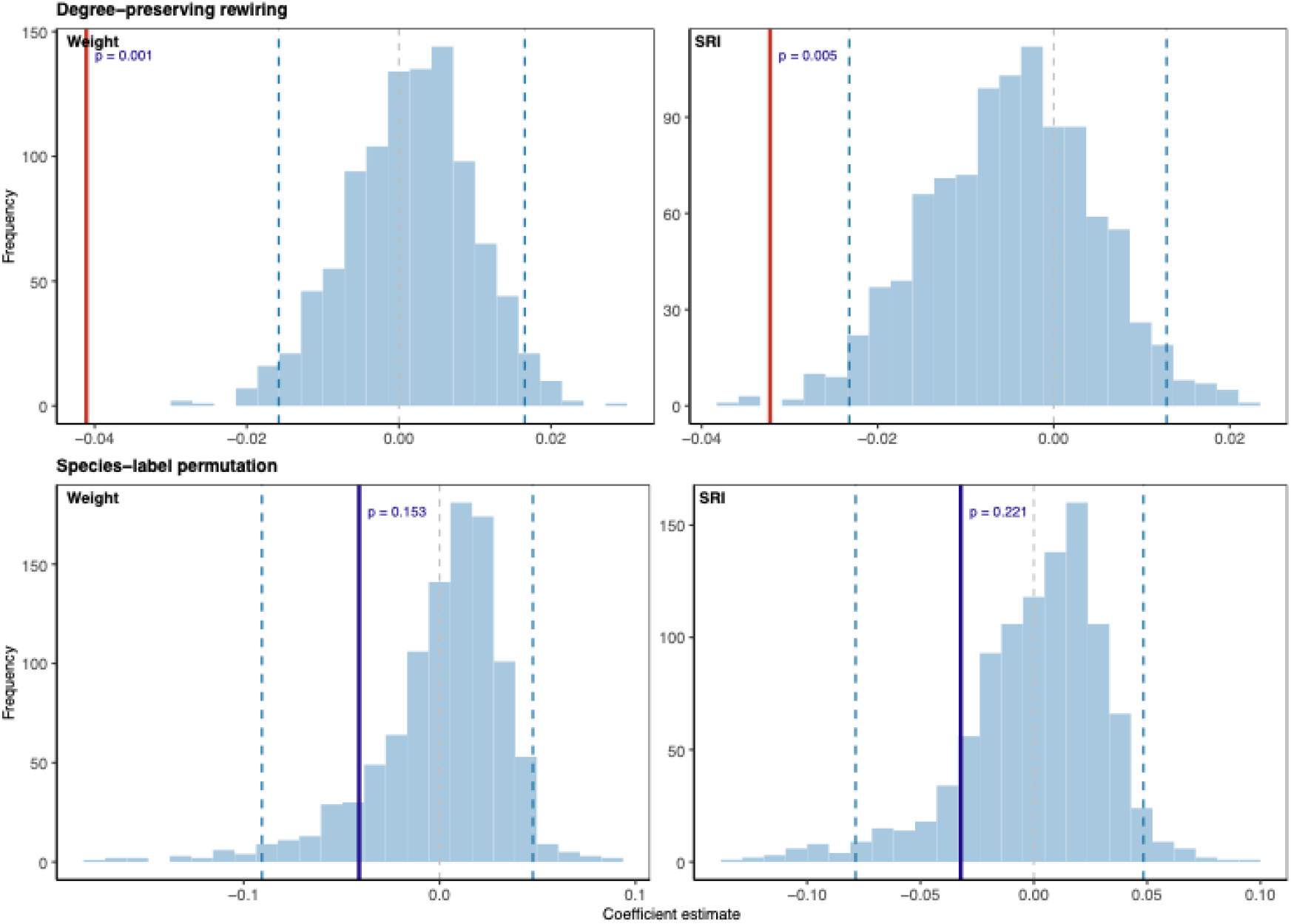
Maximum viral genetic distance. The expected distribution of permuted estimates in blue, with 95% confidence limits (blue dotted lines), plotted against the observed estimate (dark vertical line), for two measures of network structure; edge weight (a measure of co-occurrence in the landscape) and Simple Ratio Index (a measure of species association). Estimates from models using maximum genetic distance between sequence pairs as the response variable. Under degree-preserving rewiring (a), that shuffles species pairs while preserving the degree distribution, both measures fell outside the permuted distribution (Weight p = 0.001; SRI p = 0.005), with negative estimates indicating that more strongly associated species pairs carried more similar viruses. Neither measure differed from the species-label null (b), which preserves network structure and permutes host identity (Weight p = 0.153; SRI p = 0.221).

We modelled sequencing effort separately to account for the confound to maximum genetic distance. Log sequencing effort predicted maximum genetic distance (estimate = 0.64, 95% CI = 0.34 to 0.94), but including it as a fixed effect left both results unchanged (SRI: −0.033, 95% CI = −0.055 to −0.011; Weight: −0.041, 95% CI = −0.065 to −0.017), and outside the rewired null distribution (SRI p = 0.003; Weight p = 0.001).

Using mean genetic distance in place of maximum, neither predictor differed from the rewiring null (Figure 3, Figure 4; SRI: estimate = −0.021, 95% CI = −0.056 to 0.015, p = 0.130; Weight: estimate = −0.010, 95% CI = −0.047 to 0.028, p = 0.452), nor under species-label permutation (SRI p = 0.576; Weight p = 0.874). Using minimum genetic distance, both estimates were negative but neither credible interval excluded zero (Figure 4; SRI: −0.005, 95% CI = −0.056 to 0.046; Weight: −0.034, 95% CI = −0.087 to 0.021).

**Figure 3.**
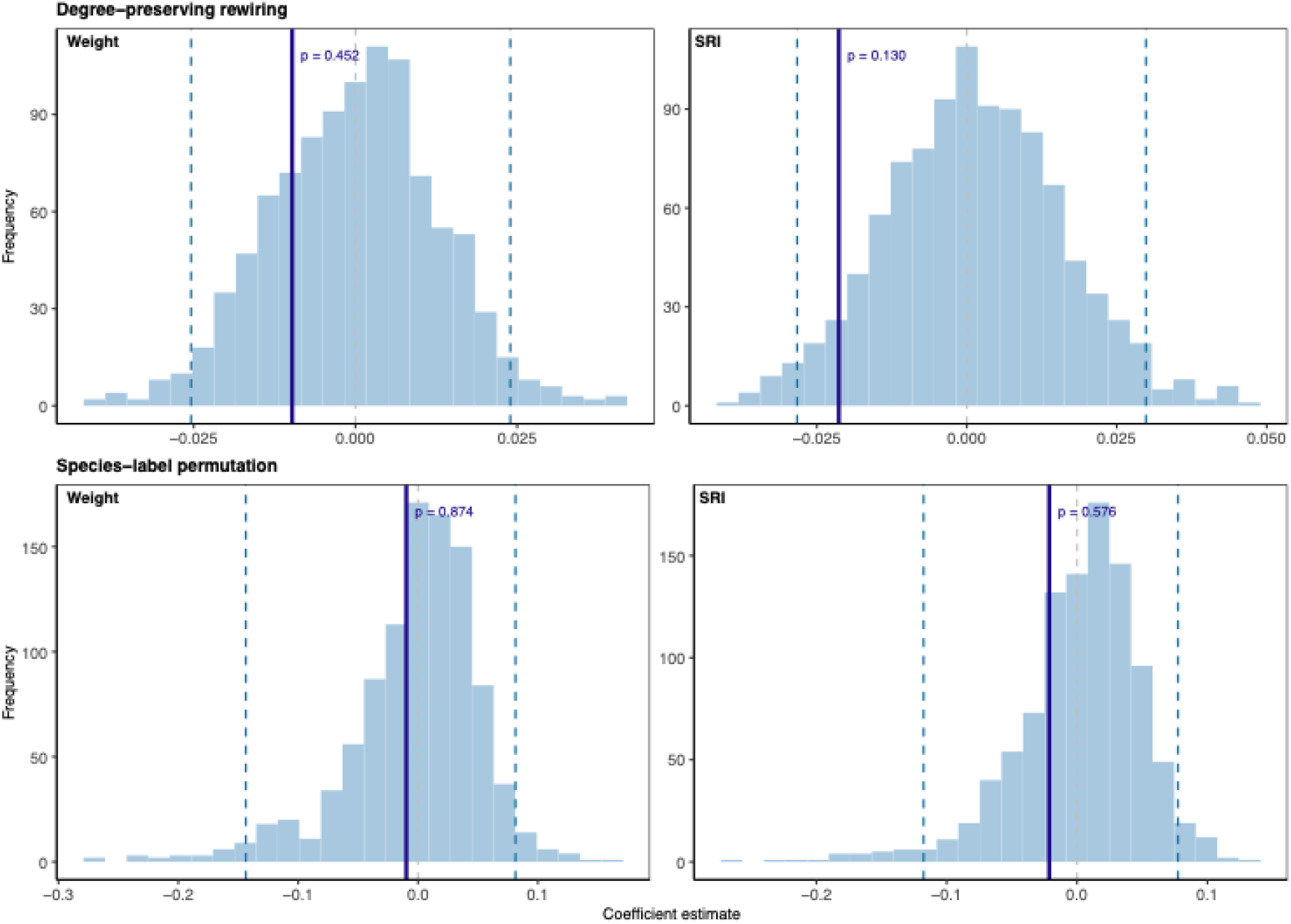
Mean viral genetic distance. As Figure 2, but for models using mean genetic distance between sequence pairs as the response variable. Neither measure differed from the permuted distribution under degree-preserving rewiring (a; Weight p = 0.452; SRI p = 0.130) or species-label permutation (b; Weight p = 0.874; SRI p = 0.576).

**Figure 4.**
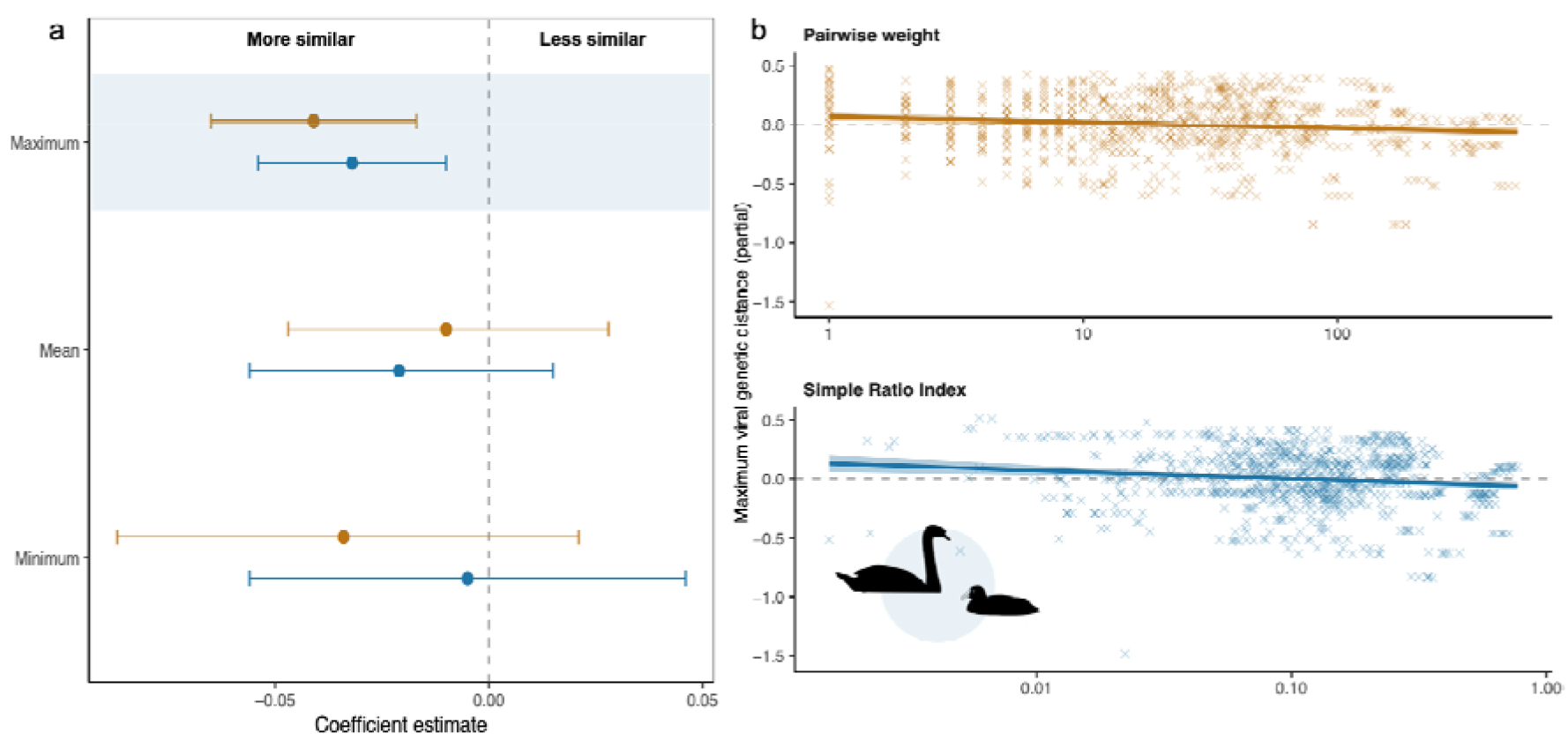
**(a)** Coefficient estimates for maximum, mean and minimum viral genetic distance, for both network measures (orange, pairwise weight; blue, Simple Ratio Index), with 95% credible intervals. Negative estimates indicate more similar viruses between more closely associated species pairs. Only maximum genetic distance gave credible intervals excluding zero (shaded), though all estimates trend negative – more similar virus. **(b)** Partial relationship between the pairwise weight and Simple Ratio Index and maximum viral genetic distance, accounting for geographic distance, temporal distance and species identity.

Temporal distance between samples strongly predicted more divergent viral genomes (Weight model: estimate = 0.196, 95% CI = 0.177 to 0.215; SRI model: estimate = 0.198, 95% CI = 0.179 to 0.217). Geometric distance between samples predicted less divergent genomes (Weight model: estimate = −0.069, 95% CI = −0.101 to −0.037; SRI model: estimate = −0.068, 95% CI = −0.101 to −0.035), but swapped direction in models fitted without the species-level random effect, reflecting substantial between-species differences in viral divergence (see Discussion). Using a multi-membership random effect structure to account for this, we found substantial variation in individual species contributions to both models (Weight model: sd = 1.42, 95% CI = 1.04 to 1.97; SRI model: sd = 1.42, 95% CI = 1.03 to 2.00), suggesting that species-level context and ecology played an important role in viral genetic distance.

## Discussion

We demonstrated that social network structure in a multi-species community of wild animals predicted genetic distance between HPAI samples, a proxy for disease transmission probability. Both raw co-occurrence frequency and relative species association predicted more similar virus, suggesting that species pairs sharing space more often carry more closely related viruses. This provides scarce empirical evidence that network edges, rather than node-level host traits, can predict pathogen spread in a complex, multi-species wild animal system.

Our results are consistent with HPAI transmission being measurable through a multi-species network of wild animals using social links as a proxy for transmission pathways. Network stability over time, which we have also demonstrated here, and the speed of survey data availability will be crucial to applying these concepts to predicting future outbreak risk (Atkinson et al. 2025). We used two large, open-data repositories, underscoring their value to research into wildlife disease surveillance (Woodward et al. 2024; Shu & McCauley 2017). Recent studies have also used similar open data to model multi-species networks (Albery et al. 2020; Van Doren et al. 2025). For example, Vickers et al. (2024) used the eBird global repository of bird survey data (Sullivan et al. 2009) to link abundance of wild birds in the landscape with HPAI incursion into poultry.

Multi-species networks of wild animals may include multiple mechanisms of transmission, and our results may be explained by direct interaction or by shared environmental exposure. Species that associate more strongly are by definition present at the same sites at the same times, and may acquire virus independently, for example from contaminated water (Giacinti et al. 2024). Our co-occurrence network design does not distinguish between these mechanisms. Our viral genome database also included samples from predatory birds with close genetic distances to colonial seabirds, and those links were likely trophic, representing ingestion of infected prey. Future work could begin to explore mechanisms of transmission using sequence-level location data matched to individual survey visits, alongside environmental sampling, and with network edges defined by mechanism.

We did not investigate the role of specific host traits in shaping putative transmission. Much previous work has examined traits such as population dynamics and host behaviour in isolation, without identifying specific transmission pathways (Vickers et al. 2024; Gamża et al. 2024; Luis et al. 2015; Olival et al. 2017; Wang et al. 2026). Instead, host traits likely drive transmission through social links, by influencing immunocompetence (Hill et al. 2016), or behaviours triggered by the presence of disease in the community (Jeglinski et al. 2024), or indirectly by shaping the social network through which disease is transferred, irrespective of the presence of disease (Albery et al. 2021; Sah et al. 2018; Kulahci & Quinn 2019; Wang 2026). It is also likely that host traits drive disease transmission at different scales. Long-distance migration has been associated with intercontinental HPAI transmission (McDuie et al. 2024; Tian et al. 2015), whereas local outbreaks in poultry have been associated with short-distance dispersal in more sedentary birds (Vickers et al. 2024; Gamża et al. 2024; Giacinti et al. 2024). Geographic distance between samples was a weak predictor in our models, and reversed direction once species identity was accounted for, consistent with an effect operating at the landscape rather than at larger, national or continental scales. Our study therefore suggests how these two scales are linked, from intercontinental incursion to outbreaks in local farmyards. Evaluation of the role of host traits in HPAI transmission may further support identification of putative bridge taxa; those critical intermediary hosts between waterfowl and poultry.

Our two network measures behaved consistently across all three viral distance metrics (Figure 4), even where neither differed from the null. We used maximum viral genetic distance as our primary measure, as a precautionary measure of relatedness. Although it was correlated strongly with sequencing effort (ρ = 0.77, against 0.43 for the mean), sequencing effort was unrelated to either network measure (ρ = 0.057 and 0.105), and including it in our models left the network estimates unchanged. Estimates using mean genetic distance were in the same direction but not distinguishable from a randomised network, and using the minimum they were smaller again with credible intervals spanning zero. Our results were also only apparent once we accounted for the species involved in our model, explaining marked differences in distribution, ecology, and how often they have been sequenced. Fitted without a species-level term, both network measures predicted that pairs that co-occurred more carried more divergent virus, possibly because widely distributed species in our network may have encountered more viral introductions. Although we used a species-label randomisation with 20 host species its permuted distribution had a standard deviation of 0.031–0.034, so non-significance may not be reliable.

In addition to the gradual accumulation of mutations, genetic distances between influenza viruses may be influenced by reassortment (Marshall et al. 2013), by which genomic segments are exchanged between viruses co-infecting the same host cell, producing larger, instantaneous increases in genetic distance. There is evidence that reassortment of recently circulating HPAI viruses has driven diversification of host specificity and seasonality of transmission (Harvey et al. 2025), contributing to the complexity of defining networks of wild hosts. For this study we used partial genetic distances between segments unaffected by reassortment. Future studies should nonetheless consider whether genotypes generated by reassortment events occupy different positions within host social networks. While we used 20,103 pairwise comparisons between 214 viral genome sequences across 20 host species, our study was limited to the 20 species for which both viral sequences and survey data were available, from 159 species recorded in the network. Proactive sampling of viruses from a more comprehensive range of species would be required to map transmission pathways with greater confidence at multiple spatial and temporal scales.

Species with high network connectivity but unsampled for HPAI (Figure 1, grey nodes) represent immediate priorities for proactive surveillance so that their HPAI genomes can be sequenced to better inform future modelling. Moreover, surveillance of HPAI has been strongly biased towards waterfowl and seabirds (Dupas et al. 2026), with far less attention paid to species that may bridge infection between water birds and livestock. Our social network approach, alongside increasing availability of open data on wild species occurrence and relative abundance, offers an opportunity for further empirical modelling to better inform surveillance and epizootic disease control policy.

## Supplementary files

S1 – Viral sequences and associated metadata: GISAID accession, sequence name, host species and order, collection date and viral genotype for every sequence contributing to the analysis (Supp1_viral_sequences.csv)

S2 – Per-species summaries: events recorded at, network degree and strength, mean and SD of dyadic weight and SRI, number of dyad-years and number of sequences per host species (Supp2_species_summary.csv)

S3 – Model data: one row per species pair per survey year, with network measures, viral genetic distances, geometric and temporal distance, and number of sequence comparisons (Supp3_model_data.csv)

S4 – Permutation results for both schemes and all three response metrics: observed estimate, null mean, SD and 95% interval, p-value, and observed against permuted row counts (Supp4_permutation_summary.csv)

S5 – Between-year network similarity: Jaccard index of shared dyads and Spearman correlation of dyadic edge weights (Supp5_network_similarity.csv)

S6 – Annual network summaries: species, events, sites and edges per survey year (Supp6_annual_networks.csv)

S7 – Annotated code files

